# Structural mechanism for amino acid-dependent Rag GTPase switching by SLC38A9

**DOI:** 10.1101/2020.07.28.225524

**Authors:** Simon A. Fromm, Rosalie E. Lawrence, James H. Hurley

**Affiliations:** Department of Molecular and Cell Biology, University of California, Berkeley and California Institute for Quantitative Biosciences, University of California, Berkeley, CA 94720, USA; Molecular Biophysics and Integrated Bioimaging Division, Lawrence Berkeley National Laboratory, Berkeley, CA 94720, USA; Howard Hughes Medical Institute and Department of Biochemistry and Biophysics, University of California at San Franscisco, CA, USA

**Keywords:** SLC38A9, Rag GTPases, mTORC1, cryo-EM

## Abstract

The mechanistic target of rapamycin complex 1 (mTORC1) couples cell growth to nutrient, energy and growth factor availability (1–3). mTORC1 is activated at the lysosomal membrane when amino acids are replete via the Rag guanosine triphosphatases (GTPases) (4–6). Rags exist in two stable states, an inactive (RagA/B^GDP^:RagC/D^GTP^) and active (RagA/B^GTP^:RagC/D^GDP^) state, during low and high cellular amino acid levels (4, 5). The lysosomal folliculin (FLCN) complex (LFC) consists of the inactive Rag dimer, the pentameric scaffold Ragulator (7, 8), and the FLCN:FNIP (FLCN-interacting protein) GTPase activating protein (GAP) complex (9), and prevents activation of the Rag dimer during amino acid starvation (10, 11). How the LFC is released upon amino acid refeeding is a major outstanding question in amino-acid dependent Rag activation. Here we show that the cytoplasmic tail of the lysosomal solute carrier family 38 member 9 (SLC38A9), a known Rag activator (12–14), destabilizes the LFC. By breaking up the LFC, SLC38A9 triggers the GAP activity of FLCN:FNIP toward RagC. We present the cryo electron microscopy (cryo-EM) structures of Rags in complex with their lysosomal anchor complex Ragulator and the cytoplasmic tail of SLC38A9 in the pre and post GTP hydrolysis state of RagC, which explain how SLC38A9 destabilizes the LFC and so promotes Rag dimer activation.

## Introduction

The Rag GTPases (Rags) recruit mTORC1 to the lysosomal membrane in response to amino acids where it is activated in response to energy and growth factor availability via Rheb (1). Unlike other small GTPases, Rags exist as obligate heterodimers of functionally redundant RagA or B (A/B) in complex with functionally redundant RagC or D (C/D). Whereas inactive Rags fail to recruit mTORC1, active Rags directly interact with the mTORC1 subunit Raptor recruiting it to the lysosomal membrane (15, 16). The lysosomal membrane protein SLC38A9 is both an amino acid transporter and an Arg-dependent Rag activator (12–14, 17, 18). The N-terminal cytoplasmic tail of SLC38A9 (residues 1-119, hereafter, SLC38A9^NT^) does not play a role in amino acid transport itself, but it is sufficient to interact with inactive Rags and stimulate mTORC1 activity (12, 13). The N-terminal tail is sequestered in the absence of bound Arg but is liberated and available to bind to Rags in its absence (19, 20), suggesting a mechanism for amino-acid dependent regulation of Rag GTPases. It has been reported that the SLC38A9 N-terminal tail regulates the Rags directly by serving as a GEF for RagA (21). However, we previously found that RagA spontaneously exchanges GDP for GTP, raising the question as to whether a GEF is actually needed in this pathway. Spontaneous nucleotide exchange by RagA is blocked, however, when the Rags are bound in the LFC (10). FLCN GAP activity, which is blocked in the LFC, is essential for the downregulation of TFE3, a member of the MiT/TFE family of transcription factors controlling lysosomal biogenesis and autophagy (10, 22–25). Thus, the LFC both blocks FLCN GAP activity and prevents spontaneous nucleotide exchange by RagA. These observations suggested the hypothesis that disassembly of the LFC and release of FLCN GAP activity is crucial for Rag activation by amino acids, particularly in respect to the mTORC1-dependent downregulation of the MiT/TFE family of transcription factors (26, 27). Here we show that the cytoplasmic tail of SLC38A9 triggers FLCN GAP activity toward RagC by destabilization of the LFC. We solved the cryo-EM structures of Rags in complex with Ragulator and SLC38A9^NT^ in the pre and post GTP hydrolysis state of RagC and discuss their implications on the current model of Rag activation.

## Results

### SLC38A9^NT^ triggers FLCN GAP activity

We reconstituted the complex comprising Ragulator, inactive Rags and SLC38A9^NT^ from purified components and tested the effect of SLC38A9^NT^ on the LFC (Fig. 1a, b). To discriminate between the nucleotides bound to RagA and RagC respectively, xanthosine-specific RagC was used throughout this study (RagC^D181N^) (8). FLCN:FNIP2 GAP activity toward RagC was enhanced in the presence of SLC38A9^NT^, consistent with a positive role in Rag activation. In the presence of SLC38A9^NT^, RagC-bound XTP was undetectable (Fig. 1c).

**Fig. 1.**
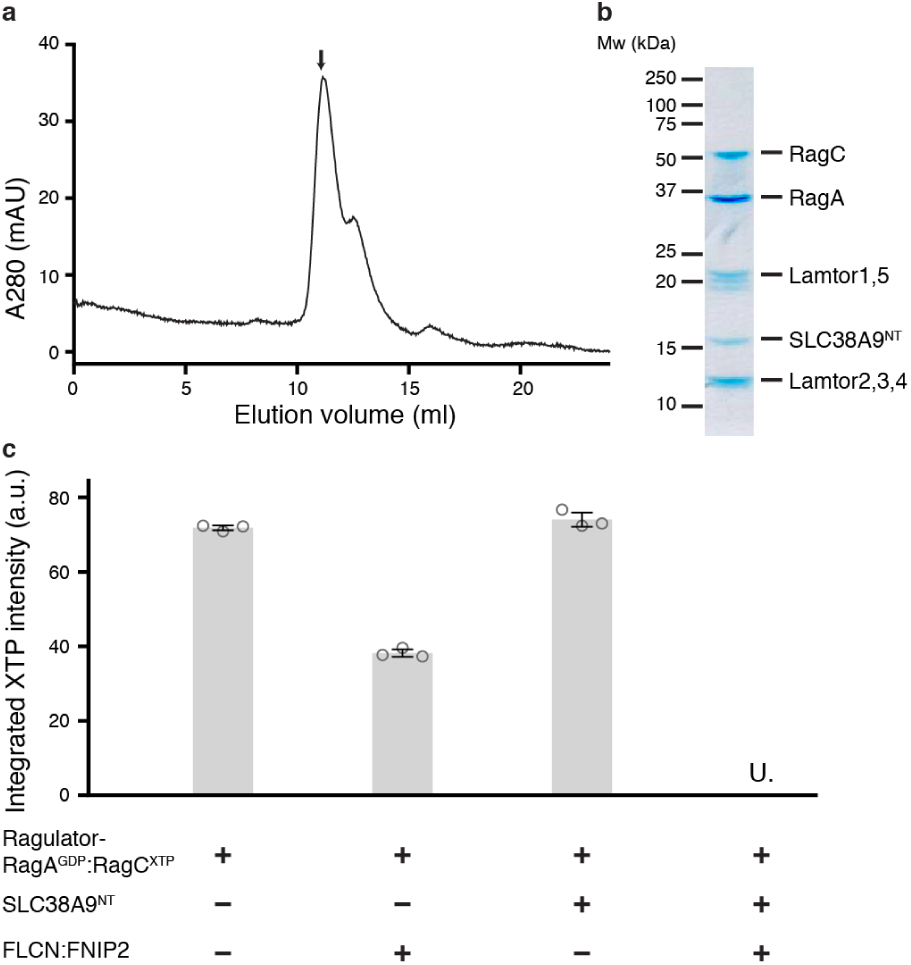
SLC38A9^NT^ triggers FLCN GAP activity. **a**, SEC profile of the reconstituted SLC38A9^NT^-RagA^GDP^:RagC^XTPγS^-Ragulator complex. **b**, SDS-PAGE analysis of the peak fraction indicated in a. **c**, HPLC-based RagC XTPase assay to measure FLCN GAP activity. Plotted are the mean ± standard deviation (*SD*) and individual data points of technical triplicates (*n*=3). U. means XTP signal was undetectable; a.u., arbitrary units.

### Structure of the pre-GAP complex

To unravel the molecular mechanism of FLCN:FNIP2 GAP activation by SLC38A9^NT^, we determined the structure of the “pre-GAP” Ragulator-RagA^GDP^:RagC^XTPγS^-SLC38A9^NT^ complex (Figs. 1a, b and 2a) by single-particle cryo-EM. The structure of this 170 kDa complex was resolved at an overall resolution of 3.2 Å (Fig. 2b and Extended Data Fig. 1). Atomic models of the Ragulator and Rag subunits were readily placed in density on the basis of previously solved structures (10, 28, 29). The remaining unassigned density was located in the cleft between the two Rag G domains and could be assigned to the central portion of SLC38A9^NT^ (residues 39-97) (Extended Data Fig. 2). The N and C-terminal portions of SLC38A9^NT^ were not resolved, consistent with previous findings that these regions are not involved in Rag binding (12). The ordered core and sole secondary structure element of SLC38A9^NT^ is an α-helical stretch deeply buried in the G domain cleft directly contacting both Rag G and roadblock domains. SLC38A9^NT^ then folds back on top of the N-core making direct contact with the GDP bound nucleotide to RagA. The most N-terminal resolved residue (Pro39) is located directly adjacent to the RagC switch I helix (α2) (Fig. 2c), whose conformation depends on the nucleotide binding state of RagC. Thus, the interaction of SLC38A9^NT^ with the Rags is intimately connected to their nucleotide binding state. SLC38A9^NT^ contains within it the so-called N-plug, which is occluded in the transmembrane domain of SLC38A9 in the absence of arginine (20). The global architecture of the Ragulator-Rag subcomplex is unchanged compared to the one observed in the LFC, but binding of SLC38A9^NT^ has two localized effects. First, it leads to a slight inward rotation of both Rag G domains and second, it occupies the cleft between the Rag G domains (Fig. 2d). Together, these effects prevent binding of FLCN:FNIP2 in the GAP-incompetent conformation observed in the LFC as multiple loops and α-helix 1 of the FLCN longin domain (α-L1) would clash with SLC38A9^NT^ and the RagA G domain, respectively (Fig. 2e). This explains how SLC38A9^NT^ binding to Rags breaks up the LFC and drives FLCN:FNIP2 into the GAP-active conformation. To gain further insight into the dynamics of the SLC38A9^NT^ interaction with inactive Rags, we measured its hydrogen-deuterium exchange (HDX) in isolation and bound to inactive Rags. In isolation, SLC38A9^NT^ shows high exchange rates (>50 %) throughout its entire sequence at the earliest time point tested, leading us to conclude the entire region is intrinsically disordered (Extended Data Fig. 3a). Consistent with the assignment from the cryo-EM structure, residues 37-97 of SLC38A9^NT^ became partially protected from H-D exchange when bound to inactive Rags (Fig. 3a, Extended Data Dataset 1). Only residues 50-55 located in a solvent exposed loop were not significantly protected at any of the exchange times (Fig. 3b and Extended Data Fig. 3b). The α-helical N-core is strongly protected, with HDX differences >30 % even after 600 s (Fig. 3a, b and Extended Data Fig. 3b, c). Thus, the HDX results corroborate the cryo-EM structure and show that the order observed for SLC38A9^NT^ is induced by formation of the complex. The SLC38A9^NT^ core harbors multiple residues shown to be crucial for the interaction with and activation of Rags in vivo (12, 13). Alanine mutations of residues Ile68, Tyr71 and Leu74, all located in the N-core, abolished the interaction with Rags (13) (Fig. 3c). Mutation of residue His60 located at the solvent exposed start of the N-core had no influence on the interaction with Rags. Mutations of stretches located in the N-plug and the part interacting with RagC α2 also abolished the Rag interaction in vivo (12) (Fig. 3c). The high consistency of the cryo-EM and HDX-MS results with the known biology of SLC38A9 mutants validates that the observed structure and dynamics correspond to the state relevant for Rag regulation in cells.

**Fig. 2.**
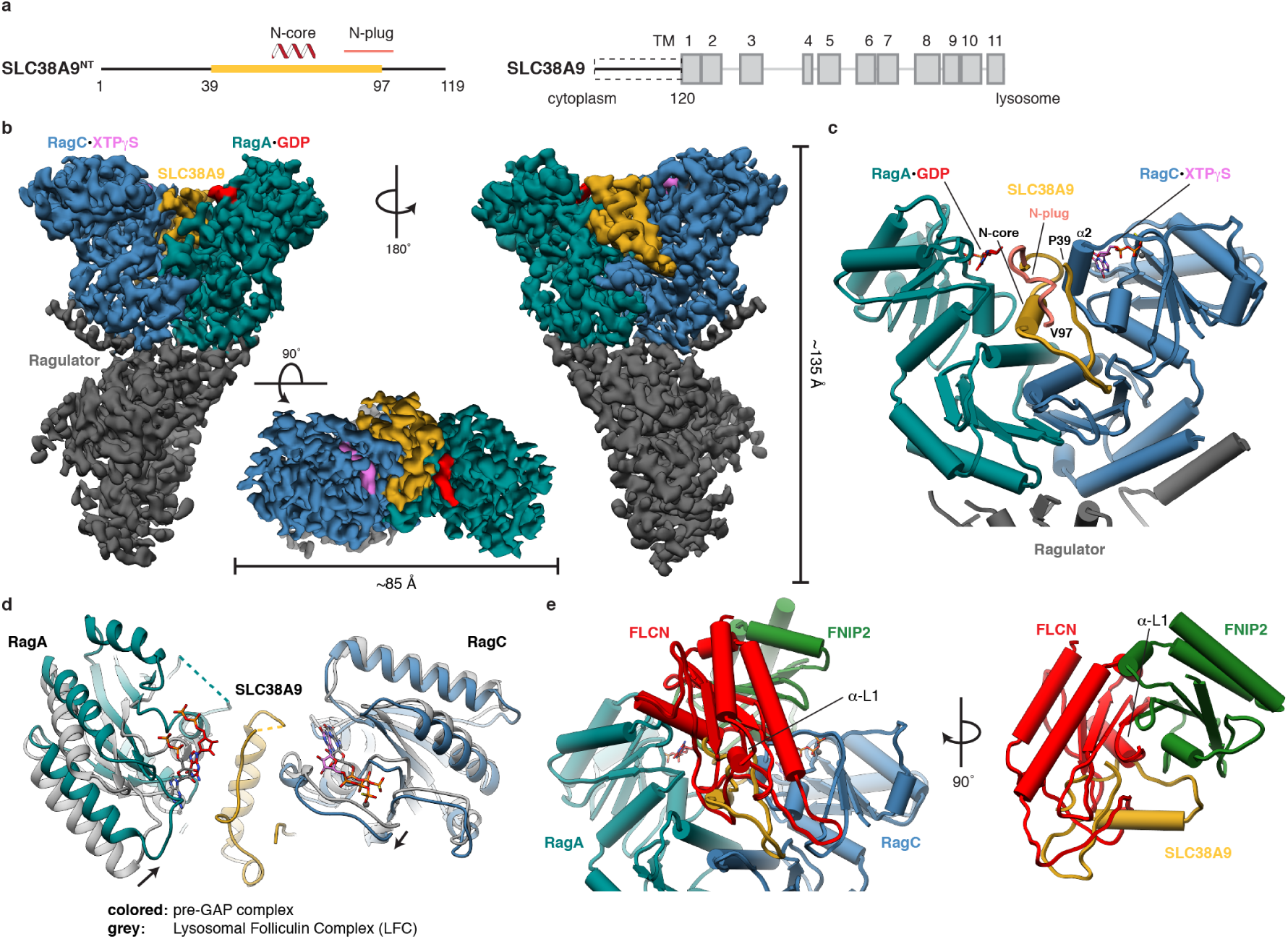
Cryo-EM structure of the pre-GAP complex. **a**, Domain organization of SLC38A9. The construct with structural annotations is shown (left, SLC38A9^NT^) in context of full-length SLC38A9 (right). Domain boundaries and cellular location of N and C-terminus are indicated. Yellow, resolved in the structure; Dashed box, used construct; TM, transmembrane helix. **b**, Cryo-EM density of the pre-GAP complex. **c**, Close-up of the SLC38A9-Rag interaction represented as pipes (α-helices) and planks (β-strands). **d**, Overlay of the pre-GAP complex (colored) with the LFC (grey). **e**, FLCN:FNIP2 (red, green) from the LFC overlaid onto the pre-GAP complex illustrating resulting clashes. Rags are omitted for clarity (right).

**Fig. 3.**
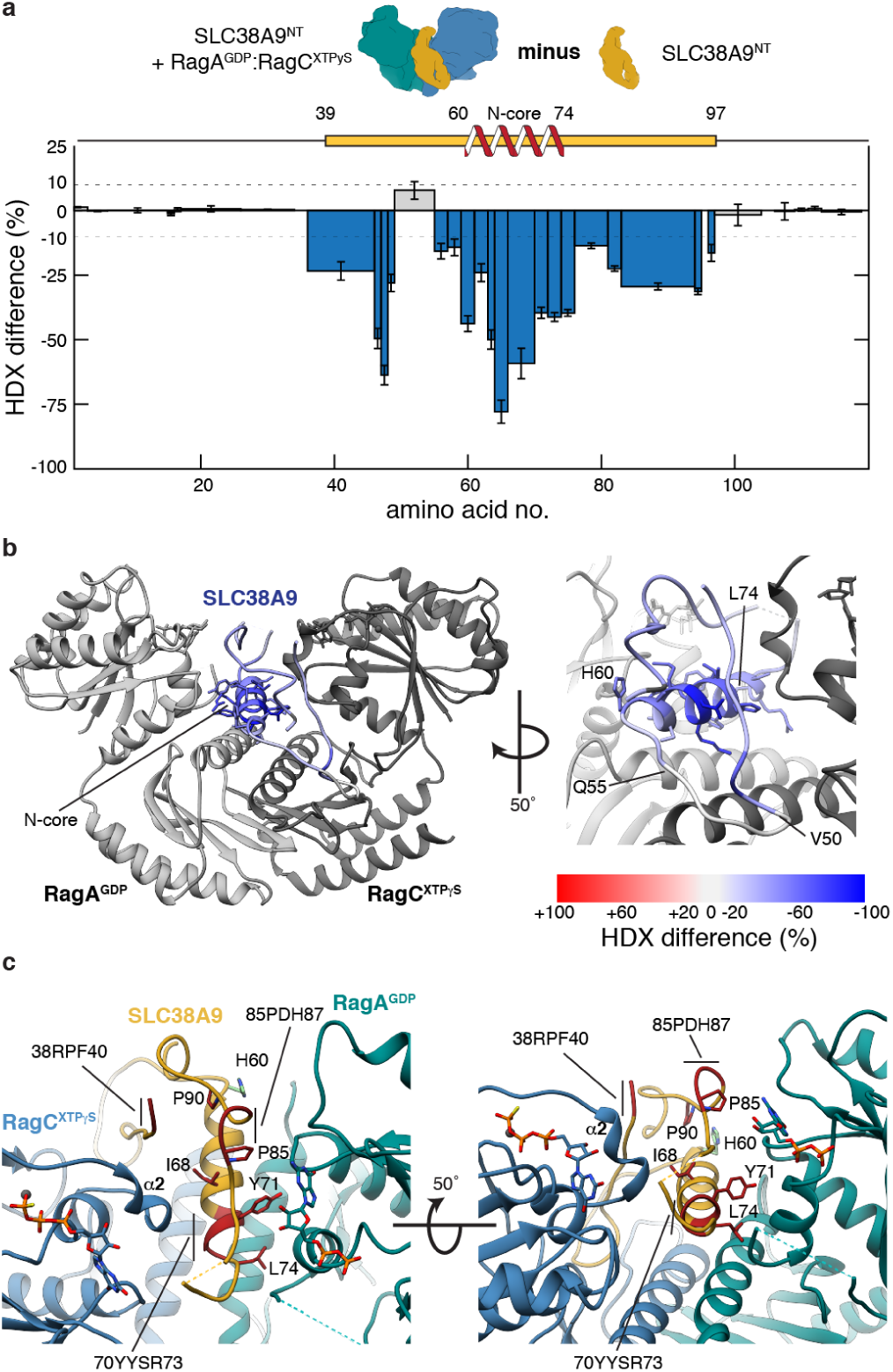
HDX-MS difference of SLC38A9^NT^ reveals complex dynamics. **a**, HDX difference plot of SLC38A9^NT^ in complex with inactive Rags and in isolation at 6 s exchange time. Plotted are the mean ± *SD* of technical replicates (*n*=4). **b**, Uptake difference mapped onto SLC38A9^NT^ in the pre-GAP complex structure. Boundaries of the N-core and the solvent exposed loop are indicated. **c**, Residues shown to be crucial for SLC38A9 function *in vivo* (red) mapped onto the pre-GAP complex structure.

### Structure of the post-GAP complex

Having shown that SLC38A9 is capable of disassembling the LFC and activating FLCN to generate the RagA^GDP^:RagC^GDP^ intermediate, we considered how this complex might progress to the active RagA^GTP^:RagC^GDP^ dimer. We found that SLC38A9^NT^, RagA^GDP^:RagC^XDP^ and Ragulator form a complex that is stable on size exclusion chromatography (Extended Data Fig. 4a, b) and will refer to it as the post-GAP complex. We used cryo-EM to solve its structure at an overall resolution of 3.9 Å (Fig. 4a and Extended Data Fig. 5). Aside from the difference in the nucleotide bound to RagC, clearly visible in density, the post-GAP complex structure is nearly identical to the pre-GAP complex structure (Cαr.m.s.d = 0.8 Å) (Fig. 4b). The RagC switch I region of the pre and post-GAP complexes are in the same conformation (Fig. 4c). In the structure of the mTORC1 subunit Raptor in complex with active Rags (15), containing RagC bound to GDP as in the post-GAP complex, the RagC switch I region was disordered and could not be resolved (Fig. 4c). In fact, all other known Rag structures bound to GDP have at least partially disordered switch I regions compared to the respective GTP bound structure (Extended Data Fig. 6). The unusual conformation is consistent with intrinsic tryptophan fluorescence data. We could not detect a FLCN:FNIP2-induced change in RagC Trp fluorescence in the presence of SLC38A9^NT^, even though GAP activity was observed by the HPLC-based GTPase assay (Fig. 1c and Extended Data Fig. 4c). Trp fluorescence reports on the conformational change of the switch I region of RagC upon GTP hydrolysis (10, 30), which the cryo-EM structure shows does not occur. Thus, the Trp fluorescence observation corroborates the unexpected finding that the double GDP-bound Rag dimer is trapped in the inactive conformation by SLC38A9^NT^. We conclude that in the post-GAP complex, RagC^GDP^ is uniquely trapped in the GTP conformation through its interaction with SLC38A9. We propose that the free energy stored by this abnormal strained conformation promotes SLC38A9 dissociation upon the appropriate trigger.

**Fig. 4.**
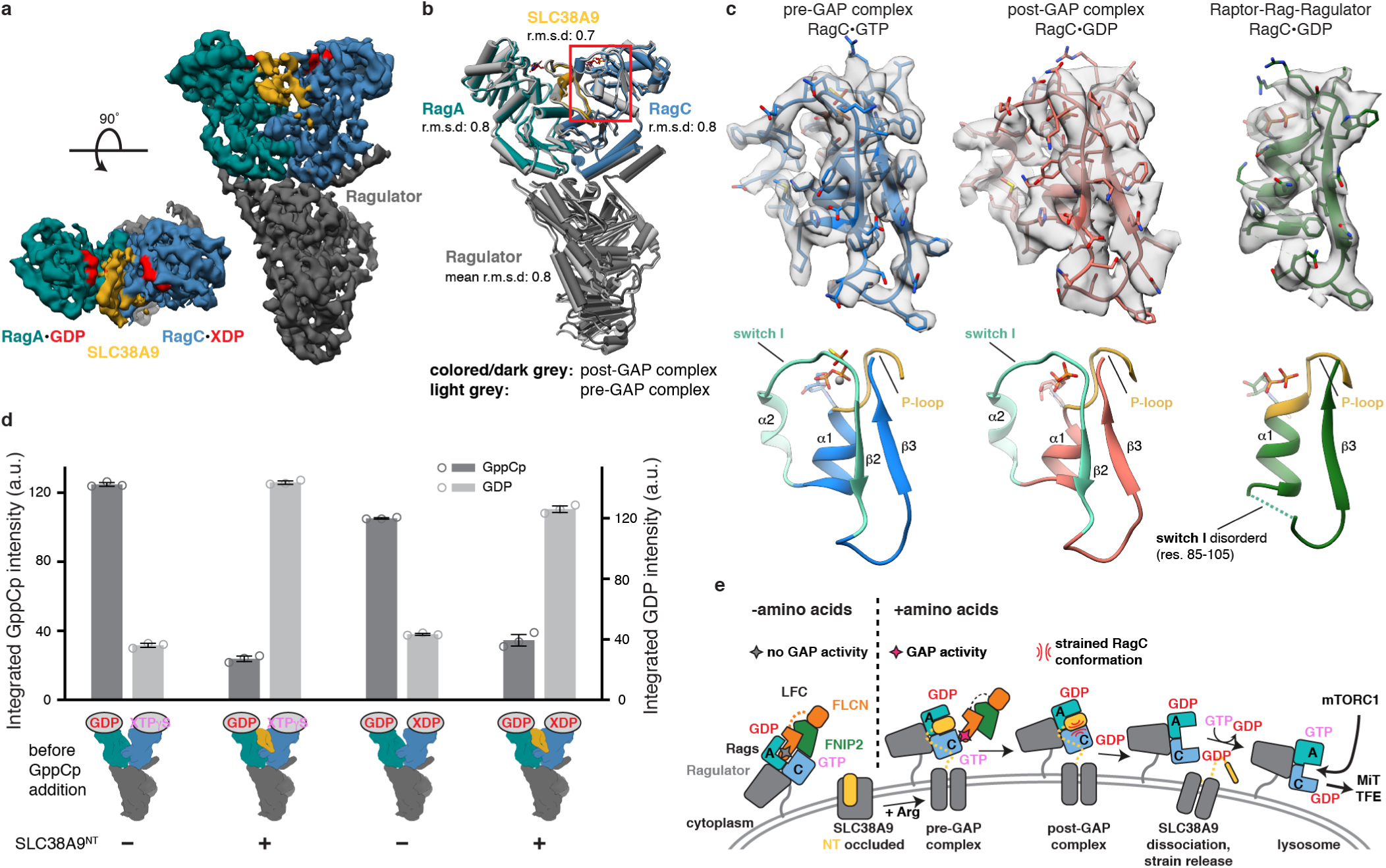
Rag GTPases are trapped in the inactive conformation in the post-GAP complex. **a**, Cryo-EM density of the post-GAP complex. **b**, Overlay of the post-GAP (dark grey, colored) and pre-GAP (light grey) complex structures represented as pipes and planks. The boxed (red) region is show in c. **c**, RagC nucleotide binding region of the pre-GAP (left, blue), post-GAP (middle, red) and Raptor-Rag-Ragulator (right, green, PDB: 6U62, EMD-20660) complex structures and respective cryo-EM density (top). Below, canonical small GTPase elements involved in nucleotide binding are highlighted in the models. **d**, HPLC-based RagA nucleotide exchange assay. Amount of GppCp bound to RagA after the assay is illustrated in dark grey, remaining GDP in light grey. Plotted are the mean ± *SD* and individual data points of technical triplicates (*n*=3). a.u., arbitrary units. **e**, Model for SLC38A9-mediated Rag GTPase activation and mTORC1 recruitment.

### SLC38A9 blocks RagA nucleotide exchange

Given that RagA spontaneously exchanges its nucleotide, that this reaction is inhibited by the LFC (10), and that the cryo-EM structures did not suggest a plausible basis for RagA GEF activity by SLC38A9^NT^, we revisited the report of RagA GEF activity by SLC38A9^NT^ (21). We employed an HPLC-based nucleotide exchange assay to directly measure GTP and GDP bound to RagA. In both pre and post-GAP complexes, spontaneous GDP-GTP exchange is greatly reduced compared to the respective Ragulator-Rag complexes in the absence of SLC38A9^NT^ (Fig. 4d). We confirmed this finding with a (2’ or 3’)-O-(N-methylanthraniloyl) (mant) GDP fluorescence-based nucleotide exchange assay (31) (Extended Data Fig. 7). These observations show that SLC38A9^NT^ is not a GEF but rather antagonizes nucleotide exchange. Whereas the HPLC and fluorescence-based assays presented here directly detect the nucleotide bound to RagA in bulk, the cross-linking assay previously interpreted as evidence for SLC38A9^NT^ GEF activity (21) only detects a sub-population of radioactively labelled nucleotides after UV crosslinking. With the structures now available, the previous cross-linking data (21) can be reinterpreted in a new light. In the pre-GAP complex, SLC38A9^NT^ directly interacts with the RagC switch I region (Extended Data Fig. 4d). The structural interaction of SLC38A9^NT^ with GDP bound to RagA would almost inevitably impact the crosslinking efficiency of GDP to RagA. Given the structures and the new biochemical data presented here, the cross-linking data (21) are best explained by changes in the accessibility of the nucleotide, rather than exchange.

## Conclusions

The cleft between the G domains of the Rag GTPase heterodimer emerges as a regulatory platform with a total of three different protein (complexes) shown to directly bind in the cleft, namely FLCN:FNIP2, SLC38A9 and mTORC1 via the Raptor subunit (10, 11, 15, 16). Due to the overlapping binding sites, these interactions cannot occur simultaneously on a Rag heterodimer and accessibility is regulated by amino acid levels via their nucleotide binding state. We propose a model where FLCN:FNIP2 binds to inactive Rag GTPases forming the LFC when amino acid levels are low. During this phase, the N-plug of the cytoplasmic tail of SLC38A9 is occluded in the arginine binding site of the SLC38A9 transmembrane domain making it unavailable for the interaction with inactive Rag GTPases (20). Upon increasing amino acid levels, arginine outcompetes N-plug binding in the transmembrane domain enabling the interaction with inactive Rag GTPases and activation of FLCN:FNIP2 GAP activity. Dissociation of SLC38A9 is facilitated by the release of conformational strain in RagC. Spontaneous nucleotide exchange on RagA ensues, generating the active Rag dimer (Fig. 4e) and switching on the downstream signaling mTORC1 to inactivate MiT/TFE-dependent transcription.

## ACKNOWLEDGEMENTS

We thank R. Zoncu for comments on the manuscript, C. Hecksel for assistance in pre-GAP complex cryo-EM data acquisition which was performed at the Stanford-SLAC Cryo-EM Center (S2C2) supported by the NIH Common Fund Transformative High Resolution Cryo-Electron Microscopy program (U24 GM129541), D. Toso, J. Remis and P. Tobias for assistance in post-GAP complex cryo-EM data acquisition. This work was supported by NIH R01GM111730 (J.H.H) and an EMBO Long-Term Fellowship (S.A.F.) and a University of California Cancer Research Coordinating Committee Predoctoral Fellowship (R.E.L.).

## AUTHOR CONTRIBUTIONS

S.A.F design and carried out all experiments and carried out all data analysis. R.E.L. performed initial GEF assays. S.A.F. and J.H.H. conceptualized the project and wrote the first draft. All authors contributed to editing the manuscript.

## DATA AVAILABILITY

EM density maps have been deposited in the EMDB with accession numbers EMD-21686 (pre-GAP complex) and EMD-21687 (post-GAP complex). Atomic coordinates have been deposited in the PDB with accession numbers 6WJ2 (pre-GAP complex) and 6WJ3 (post-GAP complex).

## Methods

### Protein expression and Purification

RagA:RagC^D181N^ GTPases (MBP tag on RagC) and Ragulator (GST tag on Lamtor1, His tag on Lamtor2) were expressed in Sf9 insect cells via baculovirus infection and purified as described in Su *et al* (29). FLCN:FNIP2 and GATOR1 were expressed in HEK293-GNTI cells and purified as described in Lawrence *et al* (10). In brief, Sf9 and HEK293-GNTI cells were resuspended and lysed in wash buffer (25 mM HEPES, 130 mM NaCl, 2.5 mM MgCl2, 2 mM EGTA, 0.5 mM TCEP, pH 7.4) supplemented with 1 % Triton X-100 and protease inhibitor (Roche, 1 tablet per 100 ml lysis buffer) by gentle rocking at 4 °C for 20 min. Lysate was cleared by centrifugation at 30,000 x g and 4 °C for 45 min. For RagA:RagCD181N, the supernatant (SN) was applied to amylose resin (NEB), washed with wash buffer (± 200 mM NaCl) and eluted by overnight (o/n) on-column TEV protease (home-made) cleavage. For Ragulator, the SN was applied to Ni-NTA resin (Thermo Scientific), washed with wash buffer (± 200 mM NaCl) and eluted with wash buffer + 250 mM imidazole. The eluate was applied to glutathione resin (GE Healthcare), washed with wash buffer (± 200 mM NaCl) and eluted by o/n on-column TEV protease cleavage. For FLCN:FNIP2 and GATOR1, the SNs were applied to glutathione resin, washed with wash buffer (± 200 mM NaCl, ± 0.1 % CHAPS) and eluted by o/n on-column TEV protease cleavage. All final eluates were concentrated using spin filters (Millipore Sigma) and purified to homogeneity via size exclusion chromatography (SEC) equilibrated with wash buffer. Codon-optimized DNA coding for SLC38A9^NT^ was cloned into a pCAG-GST vector for expression in HEK293-GNTI cells. HEK293-GNTI cells were transfected with 1 mg DNA and 3 mg PEI (Sigma-Aldrich) per liter of cells at a density of 1.3-1.8E6 cells/ml. Cells were harvested after 48-72 h, resuspended and lysed in wash buffer supplemented with 1 % Triton X-100 and protease inhibitor as described above. After clarification of the lysate by centrifugation (see above), the SN was applied to glutathione resin and incubated for 2 h at 4 °C. The resin was washed consecutively with wash buffer supplemented with 1 % Triton X-100, 1 % Triton X-100 and 200 mM NaCl and 0.1 % CHAPS, respectively. Immobilized GST-SLC38A9^NT^ was eluted from the resin by o/n TEV protease cleavage. The eluate was concentrated using spin filters and purified to homogeneity via SEC using a Superdex 75 column (GE Health-care) equilibrated with wash buffer. Purified proteins were aliquoted and flash frozen in liquid N2 and stored at −80 °C until use.

### Rag GTPase nucleotide loading

Rags were first loaded with the respective triphosphate (GTP for RagA, XTP for RagC), non-hydrolyzable triphosphate analogue (XTPγS for RagC) or mant triphosphate (mantGTP for RagA) by diluting purified Rags at least 1:10 (v/v) into calcium and magnesium-free PBS and incubating them for 10 min at room temperature (RT) in the presence of 5 mM EDTA. Then, a 10x molar excess (over Rags) of the respective nucleotides were added. After a 30 min incubation at RT, MgCl2 was added to a final concentration of 20 mM. Unbound nucleotides were removed by buffer exchange into wash buffer using a PD-10 gravity flow column (GE Health-care). Triphosphate loaded Rags were then concentrated using a spin filter to at least 1 mg/ml. If diphosphate loading (GDP or mantGDP for RagA, XDP for RagC) was desired, the respective triphosphate loaded Rags were treated with the respective GAP (GATOR1 for RagA, FLCN:FNIP2 for RagC) for 30 min at 37 °C at a ratio of 200:1 (GATOR1) or 100:1 (FLCN:FNIP2). Correct loading of Rags was confirmed by high-pressure liquid chromatography (HPLC) analysis as described below. Loaded Rags were used for experiments performed on the same day as the loading reaction.

### Pre and post-GAP complex assembly

Loaded Rags were incubated with a 1.2x molar excess of Ragulator and SLC38A9^NT^ for 30 min at 4 °C. For cryo-EM samples, the assembled complex was injected onto a Superdex 200 column (GE Healthcare) equilibrated with wash buffer. Peak fractions containing all complex components judged by SDS-PAGE analysis were pooled and concentrated using spin filters.

### Cryo-EM grid preparation and imaging

For pre-GAP complex cryo-EM, 3 µl of sample (0.25 mg/ml) were applied to freshly glow-discharged (PELCO easiGlow, 45 s in air at 20 mA and 0.4 mbar) UltrAuFoil R1.2/1.3 grids (Ted Pella). Grids were plunged into liquid ethane using an FEI Vitrobot Mark IV after 2 s blot time with the relative blot force set to −3, humidity to 100 % and temperature to 4 °C. For post-GAP complex cryo-EM, 3 µl of sample (0.5 mg/ml) were applied to freshly glow-discharged (see above) C-flat 2/1-3C-T grids (Electron Microscopy Sciences) and plunged into liquid ethane after 2 s blot time with relative blot force set to 21, humidity to 100 % and temperature to 4 °C. Data acquisition parameters are summarized in Extended Data Table 1. Data for the pre-GAP complex structure was acquired on a Titan Krios electron microscope (Thermo Fisher Scientific) operated at 300 kV, equipped with a Gatan K3 direct electron detector operating in super-resolution mode and a 100 µm objective aperture inserted. Automated movie acquisition with a 3×3 image shift pattern (9 movies per target/focus) was performed using SerialEM (32) with a nominal defocus range of −0.8 to −2.3 µm and a calibrated pixel size of 0.8522 Å. A total of 8,315 movies (59 frames per movie) were acquired with a total dose of 59 e^-^/Å^2^ over a 2.1 s exposure. Data for the post-GAP complex structure was acquired on a Talos Arctica electron microscope (Thermo Fisher Scientific) operated at 200 kV, equipped with a Gatan K3 direct electron detector operating in super-resolution mode and a 100 µm objective aperture inserted. Automated movie acquisition with cross-pattern image shift (5 movies per target/focus) was performed using SerialEM with a nominal defocus range of −1.0 to −2.5 µm and a calibrated pixel size of 1.137 Å. A total of 2,160 movies (55 frames per movie) were acquired with a total dose of 60.5 e^-^/Å^2^ over a 5.7 s exposure.

### Cryo-EM data processing

The cryo-EM data processing work-flow is depicted in Extended Data Fig. 1d. Super-resolution movies were motion and gain corrected using the Relion3 (33) MotionCor2 (34) wrapper and binned 2x by Fourier cropping. Empty micrographs or micrographs with high levels of contamination or crystalline ice were discarded during visual screening of all micrographs. Pre-GAP complex particles were picked using crYOLO (35) with the provided PhosaurusNet model (JANNI). Post-GAP complex particles were first selected reference-free from a random subset of micrographs using gautomatch-v0.53 (http://www.mrc-lmb.cam.ac.uk/kzhang/). After 2D classification in cryoSPARC v2 (36), 8 selected classes were used for template-based autopicking in Relion3. Per-micrograph CTF determination was carried out using CTFFIND4.1.13 (37). All particle extractions as well as 3D classification, per-particle CTF determination (38) and Bayesian particle polishing (39) were carried out in Relion3. All 2D classifications, ab-Initio reconstructions, heterogeneous, homogeneous and non-uniform (40) refinements were carried out in cryoSPARC v2. Throughout the data processing workflow, particles were discarded from further processing based on 2D classification (obvious ‘junk’) and 3D classification/heterogeneous refinement (incomplete/broken complexes). In a final particle selection step, heterogeneous refinement was performed against in silico generated reference maps of the complex with and without SLC38A9 bound to the cleft to enrich for particles which had SLC38A9 bound. For the pre-GAP complex, higher order aberration correction was performed in the final homogeneous refinement with one exposure group per image shift position. The final refinement run for the post-GAP structure was a non-uniform refinement. CryoSPARC v2 database files were converted into Relion star files using UCSF pyem (41) and in-house written scripts (https://github.com/simonfromm/miscEM). Density of the pre-GAP complex has been modified using Phenix ResolveCryoEM (42) followed by autosharpening as implemented in Phenix (43). Density of the post-GAP complex has been modified and sharpened locally using LocScale (44) as implemented in the CCP-EM software suite (45). All stated resolutions are according to the 0.143 cutoff of the respective gold-standard FSC (46). Density maps used for atomic co-ordinate refinement and illustration in all figures along with the respective half maps and masks used during refinement and FSC calculation have been deposited in the Electron Microscopy Data Bank (EMDB) with accession codes EMD-21686 (pre-GAP complex) and EMD-21687 (post-GAP complex).

### Atomic model building and refinement

For the pre-GAP complex coordinate model, coordinates of all five Ragulator subunits as well as RagA and RagC are based on the LFC structure (pdb 6NZD) and were rigid body fitted separately into the pre-GAP complex density map using UCSF Chimera (47). SLC38A9^NT^ was manually build de novo in Coot (48). Atomic coordinates were refined by iteratively performing Phenix real-space refinement and manual inspection and correction of the refined coordinates in Coot. To avoid overfitting, the map weight was set to 1.0 and secondary structure restraints were enabled during automated real-space refinement. In regions with low map quality, mainly located in solvent exposed parts of the Lamtor1 subunit of Ragulator, side-chain atoms were truncated to alanine (Extended Data Fig. 2c). Agreement of the final model with the experimental density was assessed by calculation of a model-map FSC (Extended Data Fig. 2a). To assess potential overfitting, a cross-validation test in which the refined model coordinates were randomly displaced by an average of 0.5 Å and re-refined against half map 1 of the final 3D reconstruction using Phenix real-space refinement with the same parameters used in the coordinate refinement described above. Model-map FSCs of the coordinates re-refined against half map 1 were calculated using the same half map (FSC work) and half map 2 (FSC test). No overfitting could be detected (Extended Data Fig. 2b). The final model was validated using Phenix comprehensive validation including MolProbity (49) and EMRinger (50) analysis. The post-GAP complex coordinate model was based on the previously refined pre-GAP coordinate model. After rigid body fitting using UCSF Chimera, coordinates were refined following the protocol described above for the pre-GAP complex with the map weight set to 0.5 during automated real-space refinement to reflect the lower resolution of the density. Model assessment and validation was performed as described for the pre-GAP complex (Extended Data Fig. 5). All model and validation parameters of the pre and post-GAP coordinate models are summarized in Extended Data Table 1. Coordinate models have been deposited in the Protein Data Bank (PDB) with accession codes 6WJ2 (pre-GAP complex) and 6WJ3 (post-GAP complex), respectively.

### Hydrogen deuterium exchange mass spectrometry

Amide hydrogen exchange was carried out at 30 °C for 6, 60, 600 and 60,000 s, respectively by diluting 5 µl of 5 µM SLC38A9^NT^ or SLC38A9^NT^-RagA^GDP^:RagC^XTPγS^ with 95 µl deuteration buffer (20 mM HEPES pD 7.4, 130 mM NaCl, 2.5 mM MgCl_2_, 0.5 mM TCEP, 92.8 % total D_2_O content). The reaction was stopped after the respective exchange time by addition of 100 µl ice cold quench buffer (0.4 M KH_2_PO_4_/H_3_PO_4_, pH 2.2) and immediately flash frozen in liquid N_2_. All exchange reactions were carried out in triplicate or quadruplicate and stored at −80 °C until HPLC-MS analysis. Undeuterated reference samples were prepared the same way using wash buffer instead of deuteration buffer. Frozen samples were thawed just prior to injection onto a chilled HPLC setup with in-line peptic digestion and eluted from a BioBasic KAPPA 5 µm Capillary HPLC column (Thermo Fisher Scientific), equilibrated in buffer A (0.05 % TFA), using a 10-90 % gradient of buffer B (0.05 % TFA, 90 % acetonitrile) over 24 min. Desalted peptides were eluted directly onto an Orbitrap Discovery mass spectrometer (Thermo Fisher Scientific) operated with a 3.4 kV spray, 37 V capillary and 120 V tube-lens voltage. The HPLC system was extensively cleaned between samples. Initial peptide identification was performed with undeuterated samples via tandem MS/MS experiments. A Proteome Discoverer 2.1 (Thermo Fisher Scientific) search was used for peptide identification and coverage analysis against entire complex components, with precursor mass tolerance ± 10 ppm and fragment mass tolerance of ± 0.6 Da. Mass analysis of the peptide centroids was performed using HDExaminer (Sierra Analytics), followed by manual verification of each peptide.

### HPLC nucleotide analysis

For analysis and quantification of nucleotides bound to Rags, 20 µl of at least 1.0 mg/ml Rags were denatured by incubation at 95 °C for 5 min and pelleted by centrifugation for 10 min at 21,000 x *g*. Released nucleotides in the SN were injected onto an HPLC system with a ZORBAX Eclipse XDB-C18 column (Agilent Technologies). Nucleotides were eluted with HPLC buffer (10 mM tetra-n-butylammonium bromide, 100 mM potassium phosphate pH 6.5, 7.5 % acetonitrile) and detected by absorbance at 260 nm. The area under the peak for quantification was calculated with the Agilent ChemStation software operating the HPLC system.

### HPLC RagC GTPase assay

The assay was carried out in a final volume of 25 µl with 13.5 µM Rags and a 1.2x molar excess of Ragulator and SLC38A9^NT^ in triplicates. The GTPase reaction was started by addition of 10 µl FLCN:FNIP2 to Rags, Ragulator and optionally SLC38A9^NT^ to yield a final FLCN:FNIP2 to Rag ratio of 1:100. Prior to addition of FLCN:FNIP2, the mixed proteins were incubated for 30 min at 4 °C to ensure complete complex formation. After addition of FLCN:FNIP2, samples were incubated for 30 min at 37 °C. Bound nucleotides after the reaction were analyzed by HPLC as described above. Plotted are the mean, standard deviation and each individual data point for each sample.

### HPLC RagA nucleotide exchange assay

The assay was carried out in a final volume of 25 µl with 14.6 µM Rags and a 1.2x molar excess of Ragulator and SLC38A9^NT^ in triplicates. Nucleotide exchange was started by the addition of 10 µl of non-hydrolyzable GTP analogue (GppCp) to Rags, Ragulator and optionally SLC38A9^NT^ to yield a final 1.2x molar excess of GppCp over Rags. Prior to addition of GppCp, the mixed proteins were incubated for 30 min at 4 °C to ensure complete complex formation. After GppCp addition, samples were incubated for 30 min at RT. Unbound nucleotides were removed after the reaction by buffer exchange into wash buffer using a Zeba spin desalting column (Thermo Fisher Scientific). Bound nucleotides were analyzed by HPLC as described above. Plotted are the mean, standard deviation and each individual data point for each sample.

### Tryptophan fluorescence RagC GTPase assay

Fluorimetry experiments were performed using a FluoroMax-4 instrument (Horiba) and a quartz cuvette compatible with magnetic stirring (Starna Cells) and a 10 mm pathlength. The Trp fluorescence signal was collected using 297 nm excitation (1.5 nm slit) and 340 nm emission (20 nm slit). Experiments were performed in wash buffer at RT with stirring. The final concentration of Rags was 350 nM. 500 µl of wash buffer were added to the cuvette and after baseline equilibration, 20 µl of a protein mixture containing Rags, 1.2x molar excess of Ragulator and with or without 1.2x molar excess of SLC38A9^NT^ were added. After signal equilibration, the assay was started by addition of 20 µl FLCN:FNIP2 to a final concentration 35 nM and the fluorescence signal was recorded in 1 s intervals for 1,800 s. The signal prior to FLCN:FNIP2 addition was used for baseline subtraction and subsequently normalized to the signal right after FLCN:FNIP2 addition. This served also as the t=0 time point.

### RagA mantGDP nucleotide exchange assay

The assay was carried out with the same instrument and cuvette as the tryptophan fluorescence RagC GTPase assay (see above). Mant fluorescence was collected using a 360 nm excitation (10 nm slit) and 440 nm emission (10 nm slit). Experiments were performed in wash buffer at RT with stirring. The final concentration of Rags was 50 nM. 500 µl of wash buffer were added to the cuvette and after baseline equilibration, 20 µl of a protein mixture containing Rags, 1.2x molar excess of Ragulator and with or without 1.2x molar excess of SLC38A9^NT^ were added. After signal equilibration, the assay was started by addition of 20 µl of GTP to a final concentration of 5 µM (100x molar excess over Rags) and fluorescence was measured in 1 s intervals for 1,000 s. The signal prior to protein mixture addition was used for baseline subtraction and subsequently normalized to the signal right after GTP addition. This served also as the t=0 time point.

**Extended Data Fig. 1.**
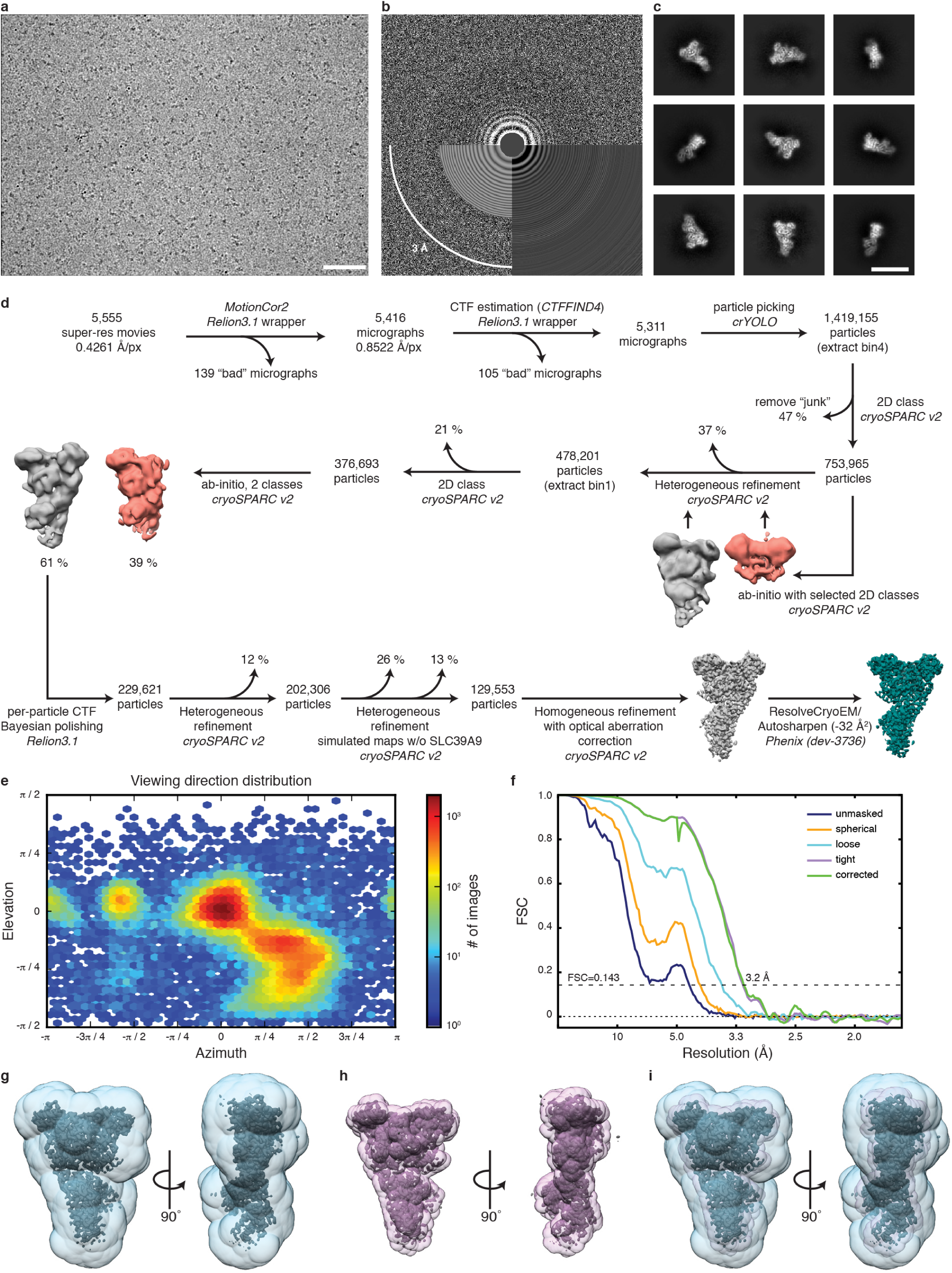
Pre-GAP complex cryo-EM structure determination. **a**, Exemplary raw cryo-EM micrograph at −2.2 µm defocus. Scale bar 50 nm. **b**, Power spectrum of micrograph shown in a with CTF estimation. **c**, Exemplary 2D class averages. Scale bar 150 Å. **d**, Cryo-EM data processing workflow. Used software is indicated with italic font. **e**, Particle orientation distribution of the final particle set. **f**, Fourier shell correlation (FSC) of the final 3D reconstruction. **g-h**, Overlay of the final density map with the masks used during refinement (g, blue, transparent), FSC calculation (h, pink, transparent). i, Both masks from g and h overlaid with the final density map.

**Extended Data Fig. 2.**
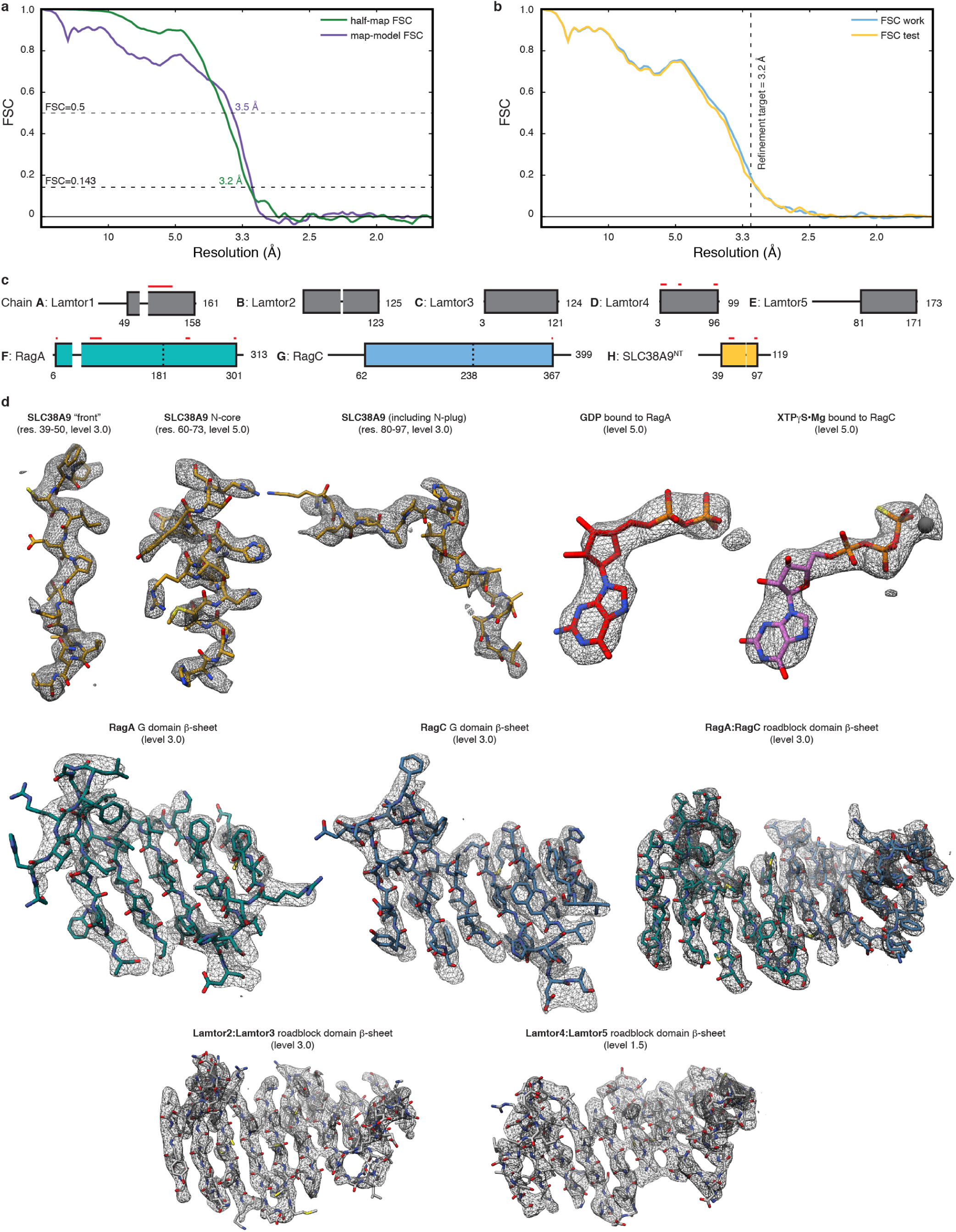
Pre-GAP complex atomic coordinate building and refinement. **a**, Overlay of half-map (green) and map-model (purple) FSC to assess map to model agreement. **b**, Overlay of FSC work (blue) and FSC test (yellow) of the cross-validation test to assess overfitting. The refinement target resolution is indicated by a vertical dashed line. **c**, Final model composition and chain assignment. Parts not resolved by the cryo-EM density are represented by thin black lines. Red lines indicate regions where side chains are truncated to alanine. **d**, Model fit in the cryo-EM density (mesh) of selected regions. The threshold level used to display the density in UCSF Chimera is given in parentheses.

**Extended Data Fig. 3.**
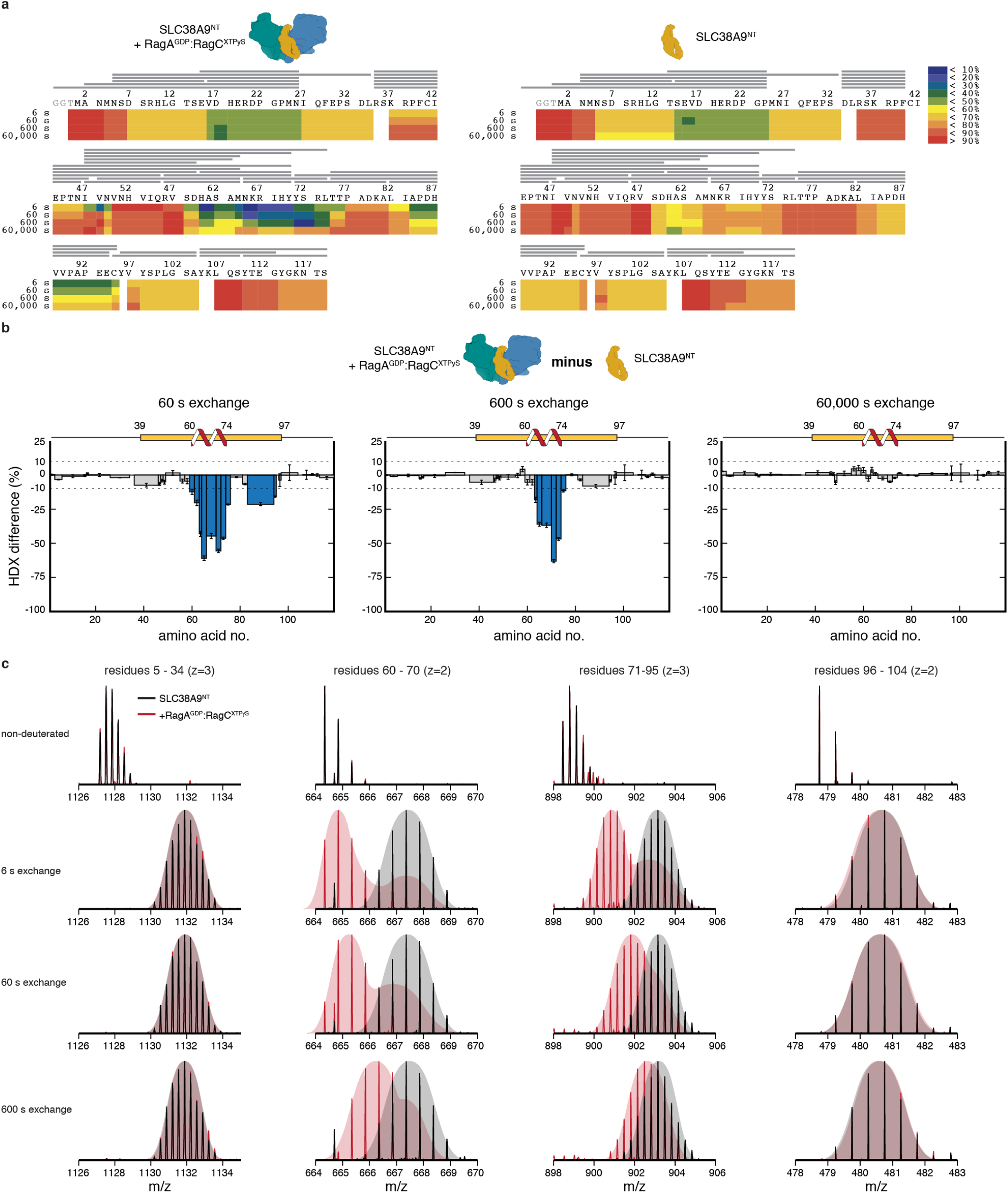
HDX-MS analysis of SLC38A9^NT^ in isolation and bound Rags. **a**, Deuterium uptake and peptide coverage (grey lines) of SLC38A9^NT^ in complex with inactive Rags (left) or in isolation (right) at 6, 60, 600 and 60,000 s exchange time. **b**, HDX difference plots of SLC38A9^NT^ in complex with inactive Rags and in isolation at 60 (left), 600 (middle) and 60,000 s (right) exchange time. Plotted are the mean +/ *SD* of technical replicates (*n*=3). **c**, Individual SLC38A9^NT^ MS peptide spectra of selected peptides in isolation (black) and in complex with inactive Rags (red). Undeuterated reference spectra are shown at the top.

**Extended Data Fig. 4.**
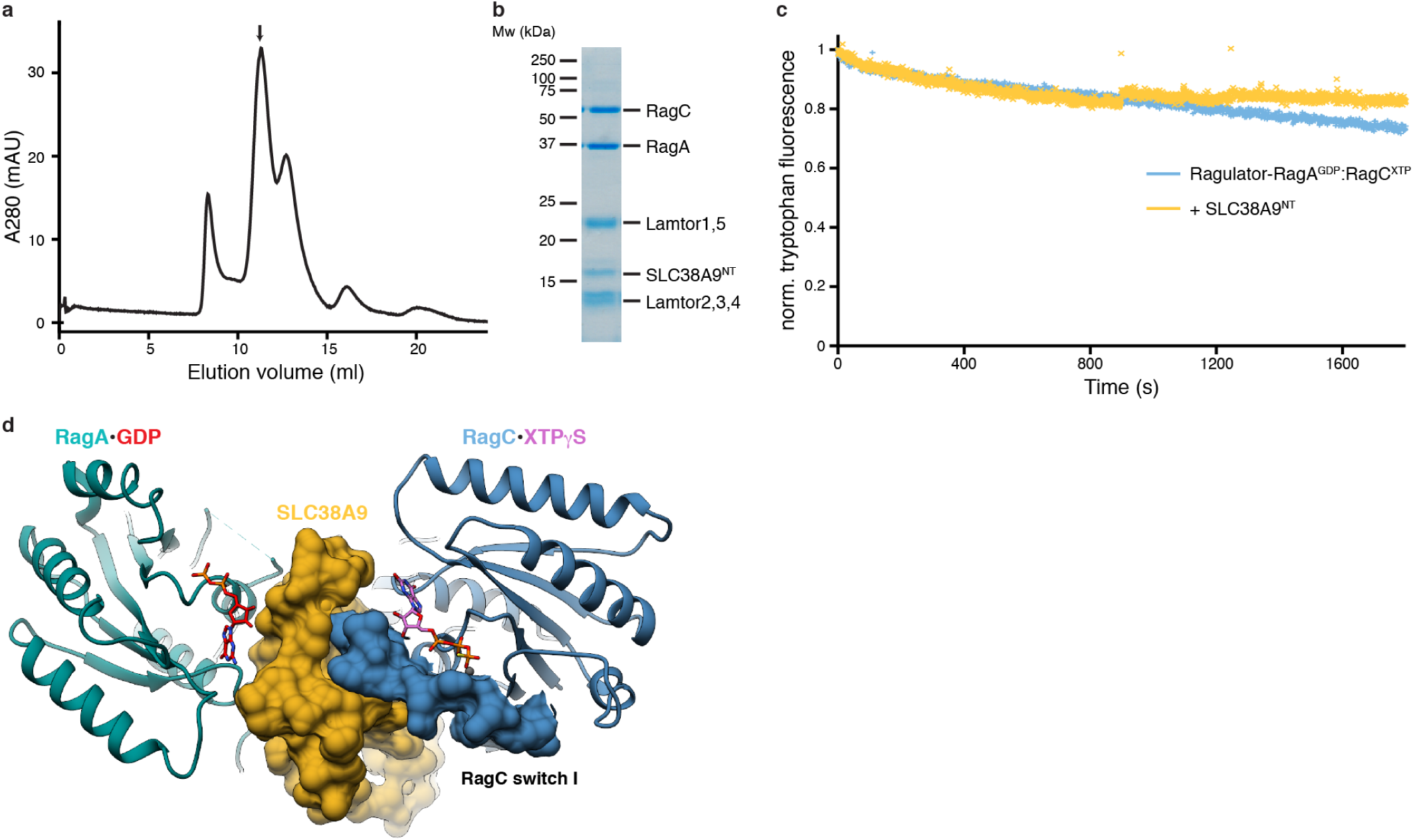
Post-GAP complex formation and Trp-fluorescence GAP assay. **a**, SEC profile of the reconstituted post-GAP complex. **b**, SDS-PAGE analysis of the peak fraction indicated in a. **c**, Intrinsic tryptophan fluorescence based RagC XTPase assay of Ragulator-RagA^GDP^:RagC^XTP^ in the absence (blue) and presence (yellow) of SLC38A9^NT^. Plotted is the normalized (norm.) Trp fluorescence of one experiment (*n*=1). **d**, Top view of the pre-GAP complex structure illustrating the SLC38A9-RagC interaction. SLC38A9 (yellow) and RagC switch I (blue) are displayes in surface representation.

**Extended Data Fig. 5.**
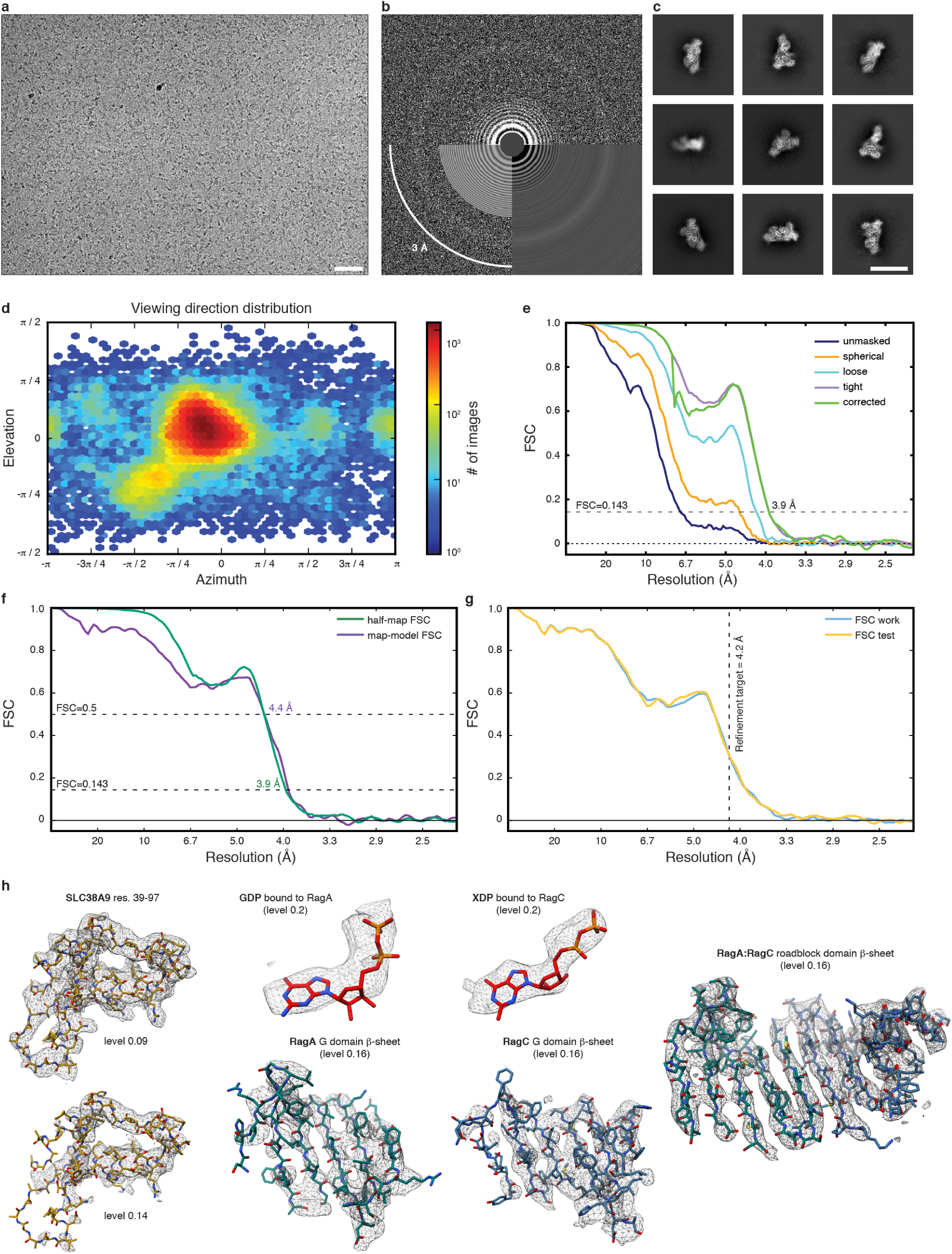
Post-GAP complex cryo-EM structure and coordinate model. **a**, Exemplary raw cryo-EM micrograph at −2.2 µm defocus. Scale bar 50 nm. **b**, Power spectrum of micrograph shown in a with CTF estimation. **c**, Exemplary 2D class averages. Scale bar 150 Å. **d**, Particle orientation distribution of the final particle set. **e**, Fourier shell correlation (FSC) of the final 3D reconstruction. **f**, Overlay of half-map (green) and map-model (purple) FSC to assess map to model agreement. **g**, Overlay of FSC work (blue) and FSC test (yellow) of the cross-validation test to assess overfitting. The refinement target resolution is indicated by a vertical dashed line. **h**, Model fit in the cryo-EM density (mesh) of selected regions. The threshold level used to display the density in UCSF Chimera is given in parentheses.

**Extended Data Fig. 6.**
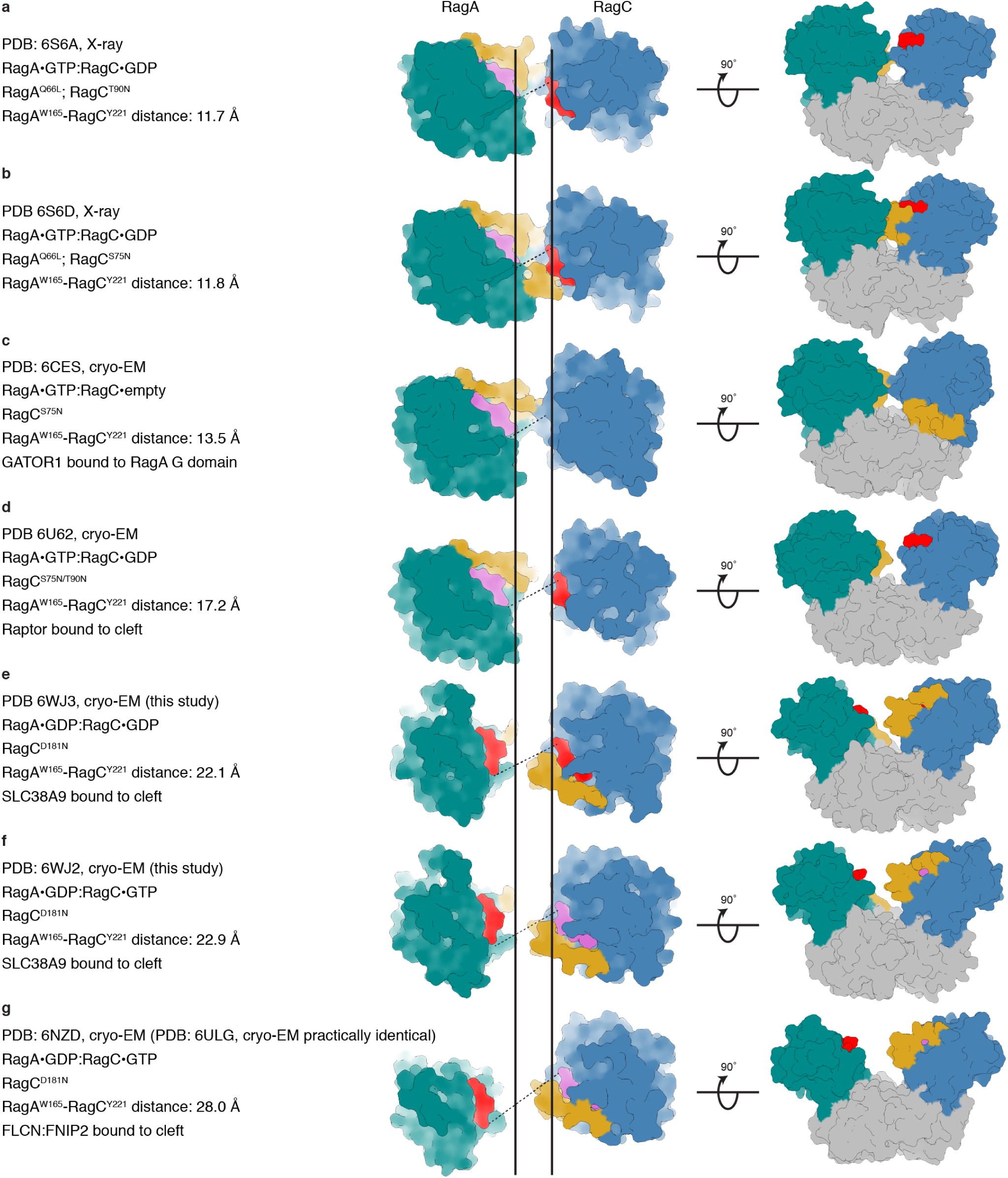
Overview of Rag GTPase structures in different states. **a-g**, Top (middle) and side view (right) of published Rag GTPase structures in surface representation (cyan, RagA G domain; blue, RagC G domain; pink, GTP or GTP analogue; red, GDP or GDP analogue; yellow, RagA or RagC switch I region; grey, RagA or RagC C-terminal roadblock domain). The dashed line connects the Cα atoms of RagA Trp165 and RagC Tyr221 representing the width of the G domain cleft. The two vertical solid lines represent the RagA Trp165 and RagC Tyr221 Cα position in a. PDB codes, experimental method, *de facto* nucleotide state and Rag binding partners (if any) are summarized on the left.

**Extended Data Fig. 7.**
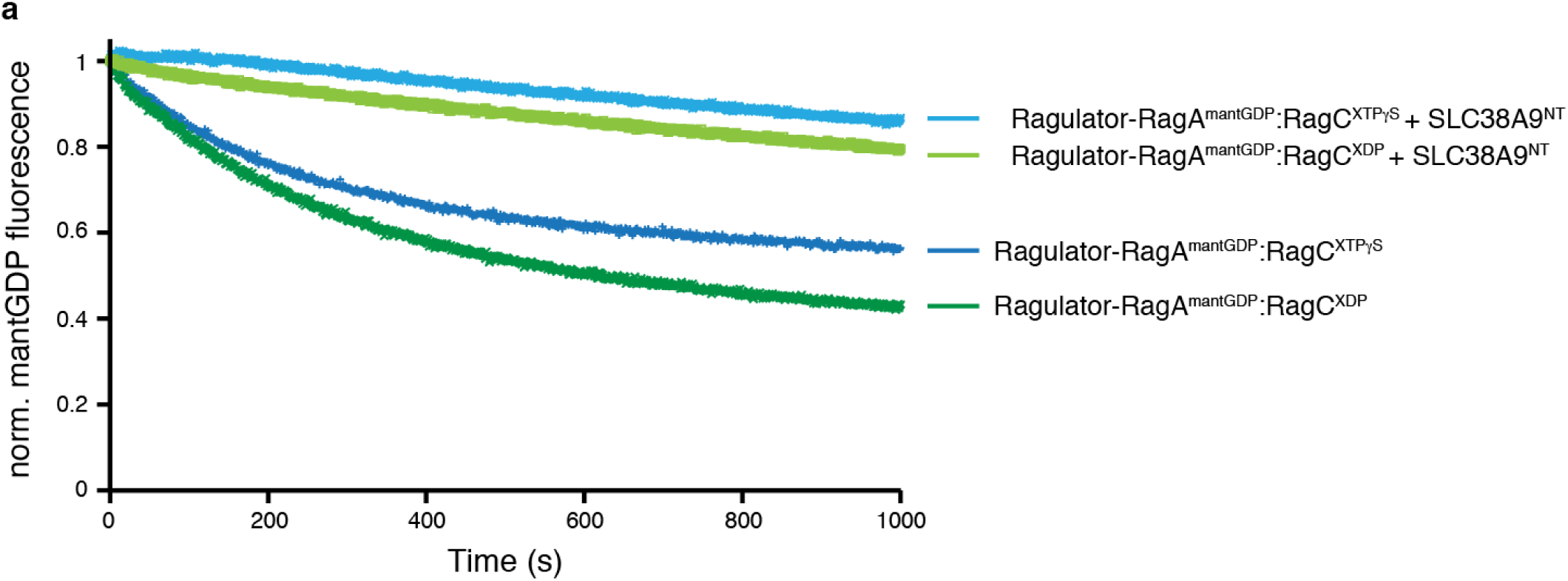
MantGDP fluorescence-based RagA nucleotide exchange assay. **a**, MantGDP fluorescence-based RagA nucleotide exchange assay of Ragulator-RagA^mantGDP^:RagC in the absence (dark blue, dark green) and presence of SLC38A9^NT^ (light, blue, light green) and RagC bound to XTPγS (light blue, dark blue) or XDP (light green, dark green), respectively. Plotted is the normalized (norm.) mantGDP fluorescence of one experiment (*n*=1).

**Extended Data Table 1:**
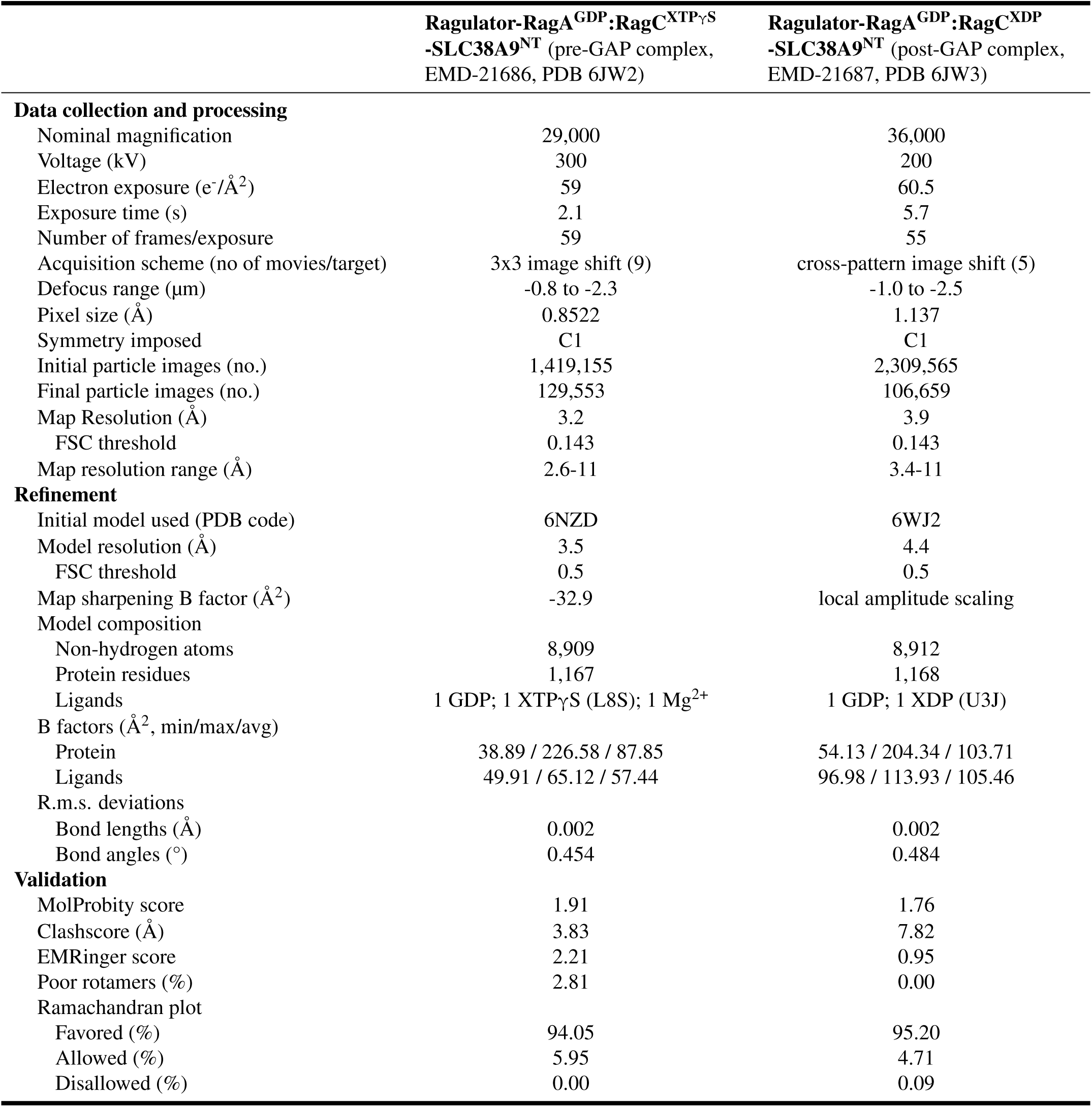
Cryo-EM data collection, refinement and validation statistics.

**Extended Data Dataset 1:**
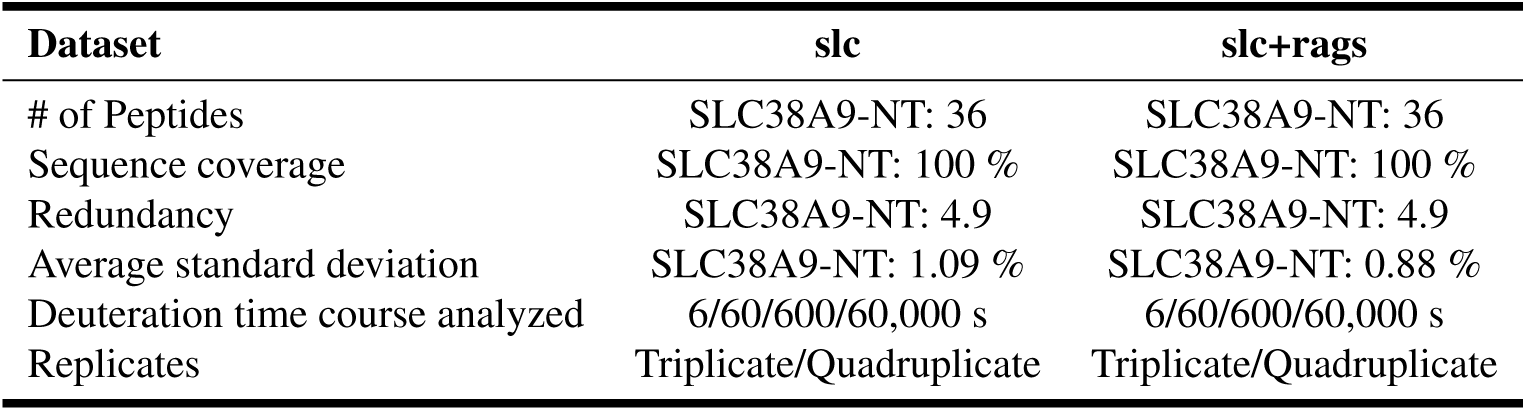
HDX-MS data summary.

